# Mechanochemical rules for shape-shifting filaments that remodel membranes

**DOI:** 10.1101/2022.05.10.490642

**Authors:** Billie Meadowcroft, Ivan Palaia, Anna-Katharina Pfitzner, Aurélien Roux, Buzz Baum, Anđela Šarić

## Abstract

The sequential exchange of filament composition to increase filament curvature was proposed as a mechanism for how ESCRT-III polymers deform and cut membranes. The relationship between the filament composition and its mechanical effect is lacking. We develop a kinetic model for the assembly of composite filaments that includes protein–membrane adhesion, filament mechanics and membrane mechanics. We identify the physical conditions for such a membrane remodelling and show this mechanism is efficient because sequential polymer assembly lowers the energetic barrier for membrane deformation.

---

Nanoscale filaments that can produce work to reshape membranes are key for cell traffic, motility, and healing. These include ESCRT-III proteins, which form polymers that reshape and cut membrane surfaces [1, 2] and are involved in a range of membrane remodelling events, including membrane budding, cell division and membrane repair [3–7]. ESCRT-III proteins come in many variants that co-polymerize to form membrane-associated composite filaments. These composite polymers dynamically exchange different monomer types with the surrounding cytoplasm. Monomer exchange is driven by the ATPase activity of an enzyme called Vps4, which extracts certain monomers while new monomers are added to the polymers that remain [2, 8, 9].

Recent experiments [10, 11] have suggested that the composition of ESCRT-III filaments changes with time in a precise sequence as monomers assemble, and are removed and replaced by monomers of a different type, in a fixed order, as shown in Fig. 1a. Additionally, although monomers of different types share a similar molecular structure [2, 5], ESCRT-III polymers have been observed to have different shapes and different membrane binding interfaces, depending on their precise monomer composition [5, 12–17]. Therefore, as the co-polymers change in composition, their geometry and mechanical properties are also expected to change, allowing ESCRT-III to perform work on the membrane [8, 18, 19]. The time-ordering of this staged assembly/disassembly process is robust and is believed to progressively drive membrane deformation and scission in a variety of biological processes [9, 17, 18, 20, 21]. For instance, in cargo trafficking, a membrane-bound ESCRT-III filament polymerises around a cargo in the shape of a flat spiral. As this flat spiral is remodelled over time into a three-dimensional helix (see Fig. 1b), the flat membrane is forced to become a tube. The subsequent exchange of monomer types is then believed to progressively increase the polymer curvature to constrict the membrane neck.

**FIG. 1.**
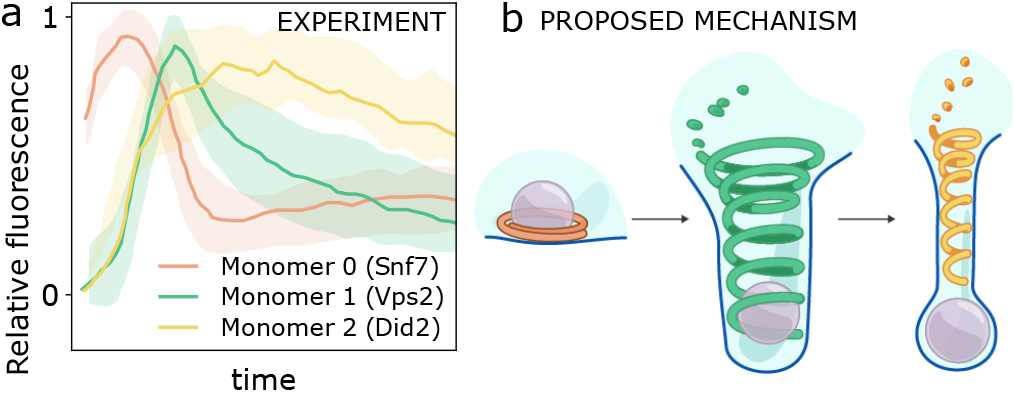
Shape-shifting by sequential monomer exchange. (a) *In vitro* experiments, tracking the presence of different monomer types (termed Snf7, Vps2, Did2) in membrane-bound yeast ESCRT-III filaments, showed that the filament composition changes according to a precise sequence [11], which is accompanied by the membrane deformation. (b) The geometry of a membrane-bound filament has been proposed to change together with its composition, driving membrane remodelling for cargo transport.

While great progress has been made on understanding this biophysical system, the physicochemical properties that enable different monomer types to assemble into dynamically shape-shifting polymers remain unclear. Here, we develop a kinetic model that can capture the recruitment and spontaneous unbinding of monomers with different physical properties. We identify rules under which the mechanical and chemical differences between different monomer types alone can drive changes in polymer assembly-disassembly kinetics and, consequently, in polymer structure. These design rules allow us to make predictions about the physiological properties of ESCRT-III monomers that are difficult to obtain experimentally, reveal the yet unexplained purpose of intermediate monomer types as catalysts for membrane deformation, and can inform efforts towards the generation of artificial shape-shifting nanomaterials.

## Model

Filaments of ESCRT-III homologs across evolution involve between ~ 3 and 11 different monomer types, as in archea [3] and animal cells [4] respectively. In addition, in systems with a large number of ESCRT-III types, some monomer types always bind together, effectively decreasing the number of assemblies [11]. To model the process, we therefore choose to use three types of monomers (*i* = 0, 1 or 2), as in Fig. 2a, which can bind to the membrane and each other, forming heterogeneous membrane-bound filaments (Fig. 2b). The filament is treated as a lattice, where each site can host up to one monomer of each type, such that different monomer types can bind side-by-side on a site. Each monomer type is associated with an energy of adhesion 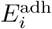, as well as a preferred curvature *c_i_*, and a preferred effective pitch *p_i_* (which is the membrane deformation depth induced by a homogeneous filament of that type). Fig. 2a shows the case in which the pure filament of type 0 is a completely flat spiral (*p*_0_ = 0) of curvature co, whereas filaments of types 1 and 2 are helices with larger pitches and curvatures, such that 0 < *p*_1_ < *p*_2_ and *c*_0_ < *c*_1_ < *c*_2_. A heterogeneous filament, made of monomers of different types, is associated with an average curvature (*C*) and pitch (*P*), which are averages of the curvatures and pitches of all its monomers.

**FIG. 2.**
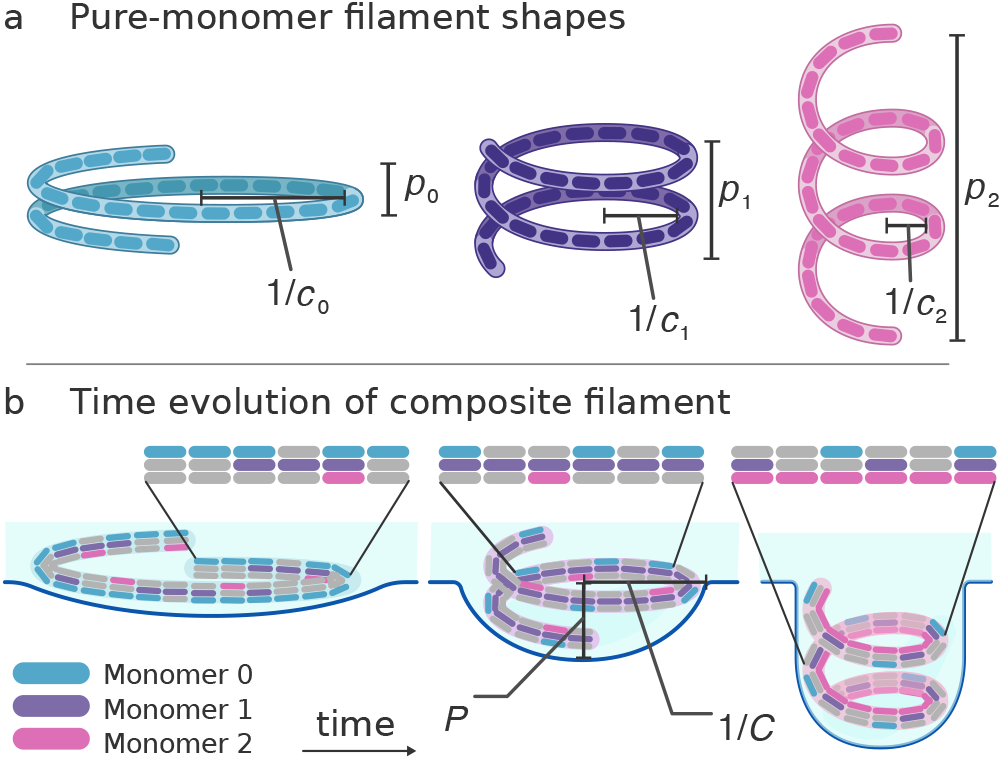
Model set up. (a) Pure filaments made up of type-*i* monomers (*i* = 0,1,2) have in-plane curvature *c_i_* and are associated with an effective pitch, *p_i_*, causing the preferred membrane deformation depth *p_i_*. (b) Three examples of the lattice composition over time with the associate overall filament shape, which gives an average pitch *P* and an average radius 1/*C*, and corresponding membrane shape. For a shallow pitch and large radius the geometry of the membrane is that of a spherical cap, while with increasing pitch or decreased radius a tube with a spherical cap end forms.

Our model considers three energy contributions: adhesion energy –*E*^adh^, filament bending energy *E*^fil^, and membrane energy *E*^mem^. The energy gained by adhesion is 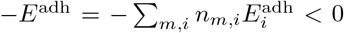 (where *n_m,i_* = 1 if site *m* contains a type-*i* monomer, 0 otherwise) and always favours monomer binding. A heterogeneous filament has a non-zero bending energy, which results from monomers of different preferred curvatures binding to the same site or to neighbouring sites, causing local elastic frustration. Assuming that sections of filament bend as elastic rods, we estimate the filament bending energy as 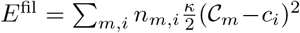 where *κ* is the filament rigidity and 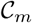 is the local filament curvature averaged over the 8 nearest neighbours of site *m*. This is based on the observation that individual monomers in ESCRT-III assemblies interact with numerous (4 to 8) other neighbouring monomers via direct contact [17, 22]. Finally, we estimate the membrane bending energy using the Helfrich model [23]: 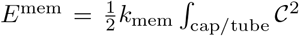 d*A* where *k*_mem_ is the membrane bending rigidity and 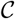 is the membrane curvature. We approximate the deformation of the membrane as a spherical cap or a cylinder with a spherical cap, thus only 2 quantities (*C* and *P*) are needed to describe the membrane shape (Fig. 2b). Further details are provided in the SI.

Overall, the adhesion term promotes binding, the filament elasticity term disfavours the neighbouring binding of monomers with different curvatures, and the membrane term penalises the binding of monomers with nonzero pitch and high curvature, which can only become part of the polymer if the membrane bends.

We investigate the dynamics of the system using the Gillespie algorithm, a variant of a dynamic Monte Carlo. Monomers bind or unbind according to rates *r*_on_ and *r*_off_. We assume that monomer concentrations in the cytoplasm are constant, as in *in vitro* experiments [11], and equal for different monomers, and that binding is diffusion-limited so that *r*_on_ is constant and sets the timescale of the simulations. The probability of a given unbinding event (i.e. a given monomer detaching from a given site) depends exponentially on the energy change associated with it: *r*_off_ = *r*_on_ *e*^*β*^(Δ*E*^adh^+*E*^fil^+Δ*E*^mem^). These energies depend on the composition of the filament, hence the unbinding rates are not just monomer-type and lattice-site dependent, but also depend on the environment of the monomer, which changes in time. We tested that all our conclusions are robust against the choice of value for *r*_on_ (Fig. S3), as long as the ratio between *r*_on_ and *r*_off_ is preserved via standard thermodynamics.

The length scale for our system is set by the reported curvature of Snf7 [12], an ESCRT-III monomer found in yeast that forms spirals of radius ~ 30 nm, which represents monomer 0 in our model. Membrane rigidity was set to 20 *k*_B_*T* [24]. Polymer rigidity is compared with indirect experimental estimates of physiological persistence lengths [13, 25]. Polymer curvatures, pitches and adhesion energies are parameters that we vary, within a range of physiologically plausible values (see table S1). Our results hold for any value of polymer length within a physiologically relevant range (Fig. S3); here we show results for a 42-site long lattice.

## Minimal design rules

In our efforts to explain the experimental data shown in Fig. 1a [11], our first task is to find the conditions that enable monomers to bind and unbind sequentially. To do so, we explored a range of values for the adhesion energies 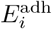, curvatures *c_i_* and pitches *p_i_* of the 3 monomer types. We quantify how much of each monomer type is present on average in the filament as a function of time, similar to the experimental fluorescence curves (Fig. 1a).

In Fig. 3a we first verify that monomers of equal properties bind together and reach an equilibrium lattice coverage, whose value is determined simply by the ratio of the binding and unbinding rates. Fig. 3b shows that sequential monomer binding emerges when monomers are assigned different pitches, irrespective of their curvatures and adhesion energies; flat monomer 0 (blue) binds before low-pitch monomer 1 (purple), which binds before high-pitch monomer 2 (pink). At time 0, the membrane is flat, so that binding of high-pitch monomers is penalised. Importantly, this initial state of the membrane creates a time asymmetry that dictates the ordering of binding. The reaction progresses until the energy of membrane deformation, induced by late-binding helix-shaped monomers, is balanced out by monomer adhesion energy. In summary, for the sequential binding to occur in the order 0 → 1 → 2, *p*_2_ > *p*_0_ > *p*_0_ must be satisfied. Although these conditions result in sequential binding, they are not sufficient to significantly deform the membrane, which requires the unbinding of the low-pitch monomers.

**FIG. 3.**
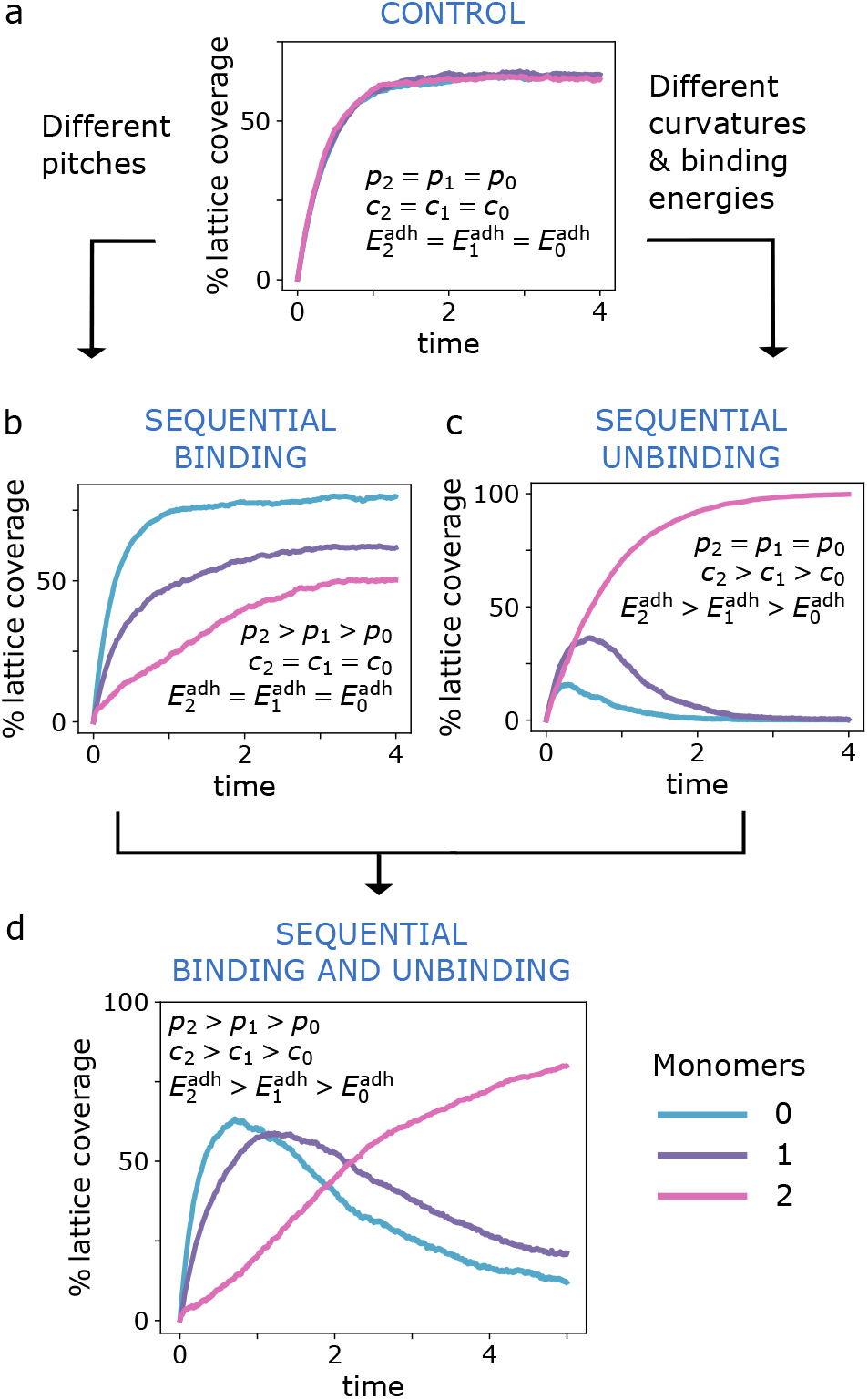
Geometry and adhesion control sequentiality. The effect of the pitches, curvatures and adhesion energies for the 3 types of monomers on the filament composition (% lattice coverage). When monomers are all equal (a) the filament stabilises as a composite mixture of the 3 monomers in equal parts. When the monomer types have different pitches (b), the binding of high-pitch monomers is delayed. Differences in curvature and adhesion energy together (c) result in the unbinding of weak-binding monomers, due to bending frustration. (d) Differences in pitch, curvature and adhesion energy are all three needed to achieve sequential assembly and disassembly. Parameters: equal pitches (a and c), *p*_0_ = *p*_2_ = *p*_2_ = 0nm; otherwise (b and d), *p*_0_ = 0nm, *p*_1_ = 30 nm, *p*_2_ = 45 nm. Equal curvatures (a and b), *c*_0_ = *c*_1_ = *c*_2_ = (15 nm)^-1^ and 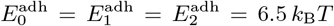; otherwise (c and d), *c*_0_ = (30nm)^-1^, *c*_1_/*c*_0_ = 2.5, *c*_2_/*c*_0_ = 8, and 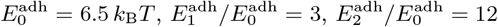. Time is in units of 10^3^ simulation steps.

Sequential unbinding in the model is a consequence of monomer frustration within a filament. If the preferred shape of a particular monomer is very different from the local average filament shape, the monomer pays a substantial cost in elastic energy and will likely spontaneously unbind. For this effect to cause asymmetric unbinding of the monomers, one monomer type must be favoured. This is the case if the monomers also have different adhesion energies, as shown in Fig. 3c. The monomer type with highest membrane-adhesion energy ends up dominating and eventually forcing the other two off the membrane. Conversely, the first monomer to unbind is the one with the smallest adhesion energy and a curvature most different from the monomer type last to be bound. Hence, for unbinding to occur in the order 0 → 1 → 2, *c*_2_ > *c*_1_ > *c*_0_ and 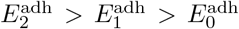 must be satisfied. The sequential disassembly fails if the monomers’ curvatures are too dissimilar, in which case prohibitive filament frustration causes monomers to unbind as soon as they bind.

Taken together, this analysis suggests that sequential subunit exchange requires increasing pitch, increasing adhesion energy and increasing curvature with time (see Fig. 3d). A failure to satisfy any one of these three criteria hinders the sequence of binding or unbinding events. The robustness of this behaviour against changes in curvature and adhesion energies is further quantified in Fig. S4.

## Sequential binding/unbinding

To explore why the sequential assembly/disassembly is needed for membrane remodelling, we next analyse different possible pathways to the final membrane-wrapped tight helix configuration. Using the parameters that result in the most pronounced sequential assembly and disassembly of the monomers (Fig. 3d), we calculate the average energy of the system as a function of monomer 2 content (Fig. 4a, turquoise curve). We then compute the same quantity by sampling configurations in which we force monomers of type 2 to bind on their own, without preceding subunits (grey curve). The binding of monomers that prefer to form tight helices (type-2) exhibits a high energy barrier, which is completely erased when the system evolves in a stepwise fashion from 0 to 1 to 2. The transient presence of the lower-curvature monomers 0 and 1 act as mechanical catalysts and deform the membrane enough to ensure the tight helical monomers can bind without having to dramatically bend the membrane from flat to highly indented in one step. This conclusion is further confirmed by two-dimensional time-dependent measurements of the energy landscape (Fig. S7). As a consequence, the sequential assembly allows the process to occur in less time (Fig. S5).

**FIG. 4.**
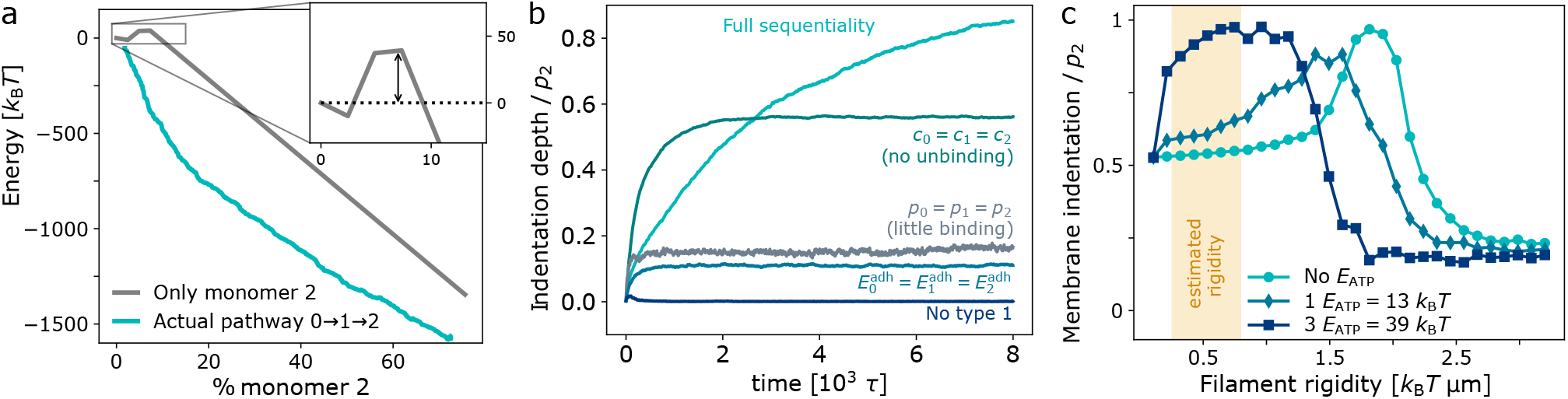
Sequentiality and membrane deformation. (a) The energy of the system as a function of type-2 monomers in the filament, when the full 0 → 1 → 2 sequence is present (turquoise), and when monomer 2 is the only one present (grey). The inset shows an energy barrier of 39 *k*_B_*T* for polymerisation of monomer 2 on membrane when no other monomers are involved. (b) Dependence of membrane deformation on monomer unbinding. If the earlier monomer types do not unbind (*c*_0_ = *c*_1_ = *c*_2_), the membrane indentation is decreased. In all the other cases (*p*_0_ = *p*_1_ = *p*_2_, 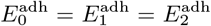), type-2 monomers do not dominate in the final filament. The largest membrane deformation is achieved with full sequential binding and unnbinding. (c) Absence of activity in our model (in turquoise), leads to sequentiality and optimal membrane indentation only for very large filament rigidities. Releasing a sufficiently large energy, 3*E*_ATP_, to elastically frustrated monomers recovers large indentation at physiologically relevant rigidities, estimated from experiments and models [13, 25] (orange band).

If the adhesion energy is high enough, tight helical monomers could in principle overcome the binding barrier and deform the membrane spontaneously. However, we expect there to be an upper limit to the adhesion energy, as in any supramolecular system, due both to the physical-chemical constraints of the membrane interaction and to the fact that active disassemblers (e.g. Vps4) need to be able to disassemble the filament to trigger neck fission using the finite energy provided by ATP hydrolysis.

Fig. 4a shows that the presence of multiple monomer types renders the binding of a highly membrane-deforming filament possible through lowering the energy barrier, but it does not explain why earlier monomer types also need to unbind. Fig. 4b shows the depth of membrane indentation for 4 different cases where monomer unbinding is suppressed. In all these cases, the final composition of the filament is different from the target one (type-2 monomers only) and, as a consequence, the extent of membrane indentation is reduced. Thus the unbinding of monomers of lower curvatures and pitches optimises the mechanical action of the filament on the membrane, and is also expected to be needed for membrane neck scission [26].

## Role of activity

All the parameters in our model are physiologically realistic (see Table S1), except for the bending rigidity of the filament. To reproduce the unbinding sequence observed in experiment, we find that our filament rigidity needs to be between 3 and 7-fold larger than previous estimates [13, 25]. This apparent mismatch is due to the fact is that our passive model does not account for active energy supply, while in experiments dynamic disassembly is only observed upon addition of ATPase Vps4, which unfolds monomers and extracts them from the filament [9, 10, 27–31]. This led us to explore whether a Vps4-like activity that promotes stress-dependent polymer disassembly could aid the process. We therefore extended the model so that a monomer with a bending stress above a certain threshold receives an additional energy penalty, a multiple of *E*_ATP_ ≃ 13*k*_B_*T*, making it more likely to unbind (details are provided in the SI). Fig. 4c shows that maximal membrane indentation can be recovered for realistic values of the filament rigidities (< 0.8 *k*_B_*T*μm) if enough energy is provided to frustrated monomers. The amount of energy needed is of the order of 3*E*_ATP_ ~ 39 *k*_B_*T*, which is consistent with the energy consumed by AAA-ATPases such as Vps4 [32]. Our findings are hence in line with the hypothesis that frustrated monomers in heterogeneous polymers could be more favourably exposed to Vps4, which in turn preferentially extracts them from the polymer [18].

## Discussion

In this work we have shown how the staged assembly and disassembly of nanoscale filaments can deform membranes. Such polymers function as a dynamic shape-shifting metamaterial: a filament changes its composition, and hence its properties, over time to perform a function that each of its parts cannot achieve when acting alone. The overall effect is a membranebound elastic structure that becomes more curved with time. Over time the recruitment of high-pitch monomers is favored, which in return facilitate the unbinding of low pitch monomers, generating an increasingly deformed membrane. The dynamics of the structure is governed by the properties of the composite monomers and by membrane mechanics. To generate a deep indentation of the membrane, subunits must have different adhesion energies, different preferred curvatures, and different preferred membrane indentations, which together lowers the energy barriers for binding.

Here, we have considered only three types of monomers. However, it is likely that the inclusion of additional monomer types (as seen in many experimental systems) can further lower the minimum membrane-adhesion energy required to drive the process to completion. Nevertheless, adding more monomer exchanges costs additional time and adds complexity. It is to be expected that there is an optimal number of subunits that minimises the overall time for membrane deformation by minimising the energy barriers and number of steps in tandem.

Within our framework, the unbinding of earlier subunits is caused by filament frustration. This frustration could be realised in different ways. It could be a result of monomers bound together having different curvatures, as shown above, or different torsion rigitidies (explored in Fig. S8) or a combination of both. In addition, there is mounting evidence that the mechanical properties of the membrane also modulate ESCRT-III function by affecting the binding dynamics of the monomers [33–36]. Our model quantitatively supports this hypothesis, as monomers are more likely to bind to a membrane that is already deformed according to the filament’s preferred morphology. Finally, these insights may inform the design of dynamic cell reshaping nanomachines, such as membrane-attached DNA-origami structures [37, 38].

## Acknowledgements

We thank T. C. T. Michaels and J. Palacci for useful discussions. We acknowledge funding by the European Union’s Horizon 2020 research and innovation programme under the Marie Skłodowska-Curie grant agreement No. 101034413 (I.P.), the Royal Society Grant No. UF160266 (A.Š.), the European Research Council (ERC) under the European Union’s Horizon 2020 research and innovation programme (Grant No. 802960) (B.M., I.P., A.Š.) and the Volkswagen foundation Life grant (B.B. and A.Š).

