## Supplementary Information for "Mechanochemical rules for shape-shifting filaments that remodel membranes"

Section I of the supplementary text contains the details of the model and methods that led us to the results in the main text. Sections II - VIII contain additional data and explorations to support the findings in the main text. In particular, in section II we develop an additional observable, Degree of Sequentiality (DoS), which allows us to systematically explore the parameter space and pinpoint the region that gives staged assembly and disassembly. In section III we check the validity of our assumptions on lattice size and kinetic rates. In section IV we develop energetic scaling arguments to explain the position of this staged assembly/disassembly region in the phase-space. In sections V, VI and VII we explore the effect of filament pitch on the binding dynamics, the role of membrane-filament versus filament-filament binding, and present further results on the energy barrier in the absence of sequential binding. In section VIII we extend the model to include a filament torsional rigidity.

#### I. MODEL AND METHODS

**The Algorithm.** In our model, monomers polymerise on a one-dimensional lattice of fixed size (Fig. 2b). Each lattice site has a state assigned to it which evolves in time depending on the binding and unbinding rates of each type of monomer, which differ throughout time and throughout the lattice. For example, a lattice site may at a certain time have a monomer of type 1 in it. Then in the next time step either 1) nothing changes 2) type 1 detaches 3) type 0 attaches or 4) type 2 attaches. In particular, a monomer of type $i$ binds to an empty site at a rate $r_{\rm on}^i$, while it unbinds from the site $m$ it occupies with a rate $r_{\rm off}^{m,i}$. The $r_{\rm on}^i$ rates can be viewed as a measure of how quickly a monomer diffuses from the cytoplasm into the vicinity of the membrane so as to have a chance to bind. They were chosen to be constant and equal for all monomer types: $r_{\rm on}^i = r_{\rm on}$. This represents the situation where all monomer types are present in excess and in the same concentration in the cytoplasm. Experiments are indeed performed at high concentrations [11] and there is evidence that these are the relevant conditions even *in vivo* [7, 33]. The instantaneous $r_{\rm off}^{m,i}$ rates are calculated using the energy difference of the system before and after the unbinding event, as explained below.

The dynamics is modelled through the Gillespie algorithm (a variant of dynamic Monte Carlo) to create the correct trajectories of states in the lattice. The system is intrinsically noisy and kept at temperature $T$. Simulations typically ran for 48000 time steps, where 1 time step corresponds to 1 $r_{\rm on}^{-1}$. The data in the results section are an average over 100 runs. The lattice has 42 sites, but the results were tested against a largest lattice length, as shown in section III. The percentage lattice coverage is calculated by summing the number of monomers of a particular type that are bound on the lattice and dividing by the length of the lattice.

**Calculating Rates and Energies.** $r_{\rm off}$, the probability per unit time for monomer type $i$ to unbind from site $m$, given that it is bound there, depends on the the total energies of the system with and without that monomer. In particular,

$$r_{\rm off}^{m,i} = r_{\rm on}^i \, e^{\beta(-E_i^{\rm adh} + \Delta E_{m,i}^{\rm fil} + \Delta E_{m,i}^{\rm mem})} \,, \tag{S1}$$

where $\beta = (k_{\rm B}T)^{-1}$ is the inverse thermal energy, $-E_i^{\rm adh}$ is the adhesion energy gain associated to a monomer of type $i$ binding to the membrane, and $\Delta E_{m,i}^{\rm fil}$ and $\Delta E_{m,i}^{\rm mem}$ are respectively the changes in filament bending energy and in membrane bending energy associated to a type-$i$ monomer unbinding from site $m$. How we calculate the energy terms is outlined below.

*Adhesion energy:*
The adhesion energy of a monomer of type $i$ to the lattice is $-E_i^{\rm adh} < 0$. This energy represents adhesion to the membrane and, in a mean-field way, to the neighbouring monomers in the filament. We neglect differences in bonds between like-monomers and different monomer types and also between side-by-side monomer bonds and along-chain monomer bonds. The total adhesion energy is the sum of the adhesion energies of all bound filaments:

$$-E^{\rm adh} = -\sum_{m=1}^{L}\sum_{i=0}^{2} n_{m,i} E_i^{\rm adh} \,, \tag{S2}$$

where $n_{m,i}$ is the number of monomers of type $i$ in site $m$ ($n_{m,i} = 0$ or 1).

*Bending energy of the filament:*
We model monomer bending akin to a small section of an elastic rod that assumes the average shape of the bound monomers around it (local curvature). The energy cost of bending for a monomer of type $i$ at site $m$ is:

$$E_{m,i}^{\rm fil} = \frac{\kappa_i}{2}(\mathcal{C}_m - c_i)^2 \,. \tag{S3}$$

Here $c_i$ and $\kappa_i$ are the preferred curvature and the bending rigidity of monomer of type $i$ respectively. $\mathcal{C}_m$ is the local curvature at site $m$. The latter is estimated by taking into account all the monomers within distance $M$ from site $m$. $\mathcal{C}_m$ is then taken to be a rigidity-weighted average of the preferred curvatures of those monomers:

$$\mathcal{C}_m = \frac{1}{M_m + 1} \sum_{n=m-M}^{m+M} \frac{\sum_{i=0}^{2} n_{m,i}\kappa_i c_i}{\sum_{i=0}^{2} n_{m,i}\kappa_i}, \tag{S4}$$

where $M_m$ is the number of occupied sites between site $m - M$ and $m + M$. An unoccupied site does not contribute to the local curvature. $M$ represents the number of monomers over which one monomer may feel a force within the polymer. Given that the ESCRT-III monomers are known to interact with up to their fourth neighbours in the polymer, we take $M = 4$. For simplicity, we assume $\kappa_i = \kappa$, as discussed at the end of this section.

The total elastic energy of the filament is the sum of the energies of all bound monomers:

$$E^{\text{fil}} = \sum_{m=1}^{L} \sum_{i=0}^{2} n_{m,i} E^{\text{fil}}_{m,i}. \tag{S5}$$

The term $\Delta E^{\text{fil}}_{m,i}$ in equation (S1) is the difference in $E^{\text{fil}}$ upon unbinding of the type-$i$ monomer in position $m$. Note that it depends not only on the particular type of monomer, but also on its local environment on the lattice, through equation (S4). This bending energy term introduces an implicit cooperativity between like-monomers, or equivalently frustration of heterogeneous filaments, meaning that monomers prefer be bound near neighbours of the same curvature.

*Bending energy of the membrane:*
To model the membrane deformation induced by a filament, we introduce a pitch $p_i$ to the monomer types (see Fig. 2a). The pitch should be understood as a measure of preferred membrane indentation for a given pure filament of type $i$. We define the average pitch $P$ of the filament as

$$P = \frac{1}{L} \sum_{m=1}^{L} \frac{\sum_{i=0}^{2} n_{m,i} w_i p_i}{\sum_{i=0}^{2} n_{m,i} w_i}, \tag{S6}$$

where $L$ is the total number of lattice sites and $w_i$ is the torsional rigidity of monomer of type $i$, which we have taken to be proportional to the bending rigidity of that type and therefore equal to each other. We also define the average curvature $C$ of the filament as

$$C = \frac{1}{L_o} \sum_{m=1}^{L} \frac{\sum_{i=0}^{2} n_{m,i}\kappa_i c_i}{\sum_{i=0}^{2} n_{m,i}\kappa_i}, \tag{S7}$$

where $L_o$ is the total number of occupied lattice sites. In our approximation, the filament produces an indentation on the membrane whose shape is governed by $P$ and $C$. In particular, for a low-pitch filament, the membrane assumes the shape of a spherical cap of depth $P$ and opening $C^{-1}$, as in Fig. S1a. As the filament shifts composition and pushes more on the membrane, precisely for $P > C^{-1}$, the membrane deforms as a tube with a spherical cap at the end, as in Fig. S1b.

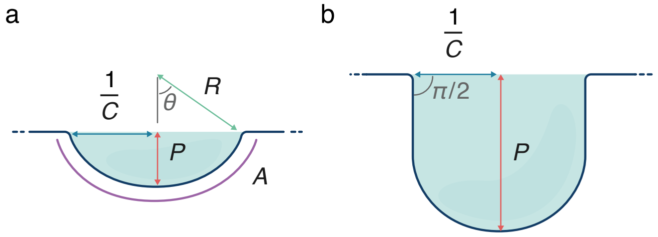


FIG. S1. **Membrane geometry.** For small pitches $P$, the membrane is a spherical cap (a). For pitches $P > 1/C$, it takes the shape of a tube closed by a hemisphere (b).

Note that there is a subtle difference between the two average quantities $C$ and $P$. The average lateral curvature only takes into account lattice sites that are occupied. If a lattice site is unoccupied, i.e. there is a space which we interpret as membrane, no curvature is added. This essentially means the membrane is not influencing the filament lateral curvature in any way. However, the average pitch is averaged over all lattice sites, occupied or not. This means the 'pitch' of an empty site is assigned as 0 and so the preferred geometry of the membrane (flat) enters the picture.

The elastic cost of the membrane bending into the shape of a spherical cap is given by

$$E^{\text{mem}}_{\text{cap}} = \frac{1}{2} k_{\text{mem}} \int_{\text{cap}} \mathcal{C}^2 \, \text{d}A, \tag{S8}$$

where $k_{\text{mem}}$ is the membrane bending rigidity and $\mathcal{C}$ is the membrane curvature. If the sphere has radius $R$, the curvature amounts to $2/R$, where the factor 2 accounts for the two principal directions of curvature. By geometry, referring to Fig. S1a, $R = (C^{-2} + P^2)/(2P)$. The area of the spherical cap is $A = 2\pi R^2(1 - \cos\theta) = \pi(C^{-2} + P^2)$, with $\theta$, $R$, $C$ and $P$ again represented in the figure. Substituting this into equation (S8), we obtain

$$E^{\text{mem}}_{\text{cap}} = 8\pi k_{\text{mem}} \frac{P^2}{C^{-2} + P^2}. \tag{S9}$$

Equation (S9) gives the expression of $E^{\text{mem}}$ when the indentation is not deep enough to produce a tube, i.e. $P < C^{-1}$.

As $P$ increases, the indentation becomes deeper and at $P = C^{-1}$ the spherical shell is a hemisphere of radius $C^{-1}$. The energy of this hemispherical cap is obtained from equation (S9) and it is independent of $C$ and $P$, as it should be:

$$E^{\text{mem}}_{\text{hs}} = 4\pi k_{\text{mem}}. \tag{S10}$$

For $P > C^{-1}$, we assume that a tube forms above the hemisphere, as in Fig. S1b. The elastic cost to produce such tube is

$$E^{\text{mem}}_{\text{tube}} = \frac{1}{2} k_{\text{mem}} \int_{\text{tube}} \mathcal{C}^2 \, \text{d}A. \tag{S11}$$

One of the principal curvatures of the tube is 0, so $\mathcal{C} = C$. The length of the tube is $P - C^{-1}$, so its area is $A = 2\pi C^{-1}(P - C^{-1})$. Substituting in equation (S11), we obtain

$$E^{\mathrm{mem}}_{\mathrm{tube}} = \pi k_{\mathrm{mem}}(PC - 1)\,. \tag{S12}$$

The membrane energy for $P > C^{-1}$ is $E^{\mathrm{mem}} = E^{\mathrm{mem}}_{\mathrm{hs}} + E^{\mathrm{mem}}_{\mathrm{tube}}$, as given by equations (S10) and (S12).

**Active energy supply.** In an extension to the model, which is used to produce the results in Fig 4c, we include an additional energy supply. This has the effect of aiding disassembly. We model this activity by adding an additional constant term, $n\,E_{\mathrm{ATP}}$, in the exponent of equation S1, where $1\,E_{\mathrm{ATP}} = 13\,k_{\mathrm{B}}T$. The energy term is added conditionally to monomers which have a bending energy term (equation S3) above a certain threshold, $E_{\mathrm{th}}$, so that the activity is stress-dependent. For Fig 4c, where strong membrane indentation is recovered for physiological filament rigidities, we used $E_{\mathrm{th}} = 3.6\,k_{\mathrm{B}}T$ and $n = 3$.

**Parameters.** We set our length scale for the system from the reported values of the radius of curvature of Snf7. This was fixed throughout the exploration at 30 nm [12]. The value for membrane rigidity set the energy scale of the system and was kept at 20 $k_{\mathrm{B}}T$ [24]. The curvatures, pitches and binding energies were then explored to find the parameters that gave sequential assembly and disassembly of types 0 and 1, with type 2 fixating at long times. This search was constrained from the discovery that differences in adhesion energy and curvature alone gave sequential unbinding, and this was independent of differences in pitch which gave sequential binding (see Fig. 3a-d). Once a set of parameters was found that gave qualitatively similar binding curves to those seen in experiment, the robustness of these parameters was explored, as described in the following sections.

TABLE S1. Model parameters

| Parameter | Model value | Literature values | Reference |
|---|---|---|---|
| Membrane rigidity | $20\,k_{\mathrm{B}}T$ | $20\,k_{\mathrm{B}}T$ | Dimova *et al.*, 2014 [24] |
| Filament* persistence length | 1.92 µm | 0.26 – 0.80 µm | Chiaruttini *et al.*, 2015 [25]<br>Shen *et al.*, 2014 [13] |
| Length per monomer in filament | 3 nm | 3 nm | Nguyen *et al.*,. 2020 [17] |
| Curvatures | $c_0^{-1} = 30$ nm<br>$c_1^{-1} = 12$ nm<br>$c_2^{-1} = 3.75$ nm | 30 nm<br>10 nm**<br>6.25 nm*** | Henne *et al.*, 2012 [12]<br>Filseck *et al.*, 2020 [15]<br>Pfitzner *et al.*, 2020 [11] |
| Adhesion energy per monomer | $E_0^{\mathrm{adh}} = -6.5\,k_{\mathrm{B}}T$<br>$E_1^{\mathrm{adh}} = -20\,k_{\mathrm{B}}T$<br>$E_2^{\mathrm{adh}} = -76.0\,k_{\mathrm{B}}T$ | $-4.0\,k_{\mathrm{B}}T$<br>$-19.0\,k_{\mathrm{B}}T$ | Chiaruttini *et al.*, 2015 [25]<br>Filseck *et al.*, 2020 [15] |
| Preferential depth of filament | $p_0 = 0$ nm<br>$p_1 = 30$ nm<br>$p_2 = 45$ nm | | |

* Persistence length, $\mathrm{L_P} = \kappa/(k_{\mathrm{B}}T)$ where $\kappa$ is the filament rigidity in equation (S3). ** Taken from an average of Filseck 'ribons and zigzag' diameters of Snf7-Vps2-Vps24 filament and assuming that the Vps2-Vps24 copolymer and Snf7 polymer contribute equally to the average diameter. *** From the radius of the membrane tube when coated with Did2-Ist1

Differences in the rigidity of the filaments were found to not make a large difference on their own to the results, and just exaggerated the effect of curvature differences. Therefore, for simplicity, the filament rigidities were kept constant and the same as each other throughout ($\kappa_0 = \kappa_1 = \kappa_2$). Table I shows the full set of parameters used (for the successful sequential peaks), together with a comparison to previously estimated values, either via experiment or other models. Rough estimates for what the monomers of type 0, 1 and 2 correspond to are 0 = Snf7, 1 = Vps2-Vps24, 2 = Did2-Ist1.

### ADDITIONAL DATA

### II. ORDER PARAMETER: DEGREE OF SEQUENTIALITY

To probe the ability of a given set of parameters to produce sequential assembly-disassembly behaviour, we construct an observable that we name Degree of Sequentiality (DoS). It is an empirical measure of how sequential the curves are, where four criteria are taken into account, described in equation (S13). The variables used for the DoS are defined in Fig. S2: $P_i(\mathrm{end})$ is the proportion of sites containing a monomer of type $i$ at the end of the simulation, $P_i^{\max} = P_i(t_i^{\max})$ is the peak of such proportion curve, and $t_i^{\max}$ is the time at which the peak is reached. The factors in front of the four terms in the DoS equation (S13) (4, 2, 1, 1 respectively) were chosen empirically so that the highest DoS corresponded phenomenologically to the most sequential-looking curves.

$$\begin{aligned}
\frac{1}{\mathrm{DoS}} \propto\ & \underbrace{4\left(\frac{P_0(\mathrm{end})}{P_0^{\max}} + \frac{P_1(\mathrm{end})}{P_1^{\max}}\right)}_{\text{are there peaks in subunits 0 and 1?}} \\
& + \underbrace{2\left|2 - \frac{P_0^{\max}}{P_2^{\max}} - \frac{P_1^{\max}}{P_2^{\max}}\right|}_{\text{are the heights of the peaks comparable?}} + \underbrace{\left(1 - \frac{P_2(\mathrm{end})}{P_2^{\max}}\right)}_{\text{is subunit 2 bound at end?}} \\
& + \underbrace{\left(\frac{P_1(t_0^{\max})}{P_0^{\max}} + \frac{P_2(t_0^{\max})}{P_0^{\max}} + \frac{P_0(t_1^{\max})}{P_1^{\max}} + \frac{P_2(t_1^{\max})}{P_1^{\max}}\right)}_{\text{are the peaks in subunits 0 and 1 high compared to the other curves at that time?}}
\end{aligned} \tag{S13}$$

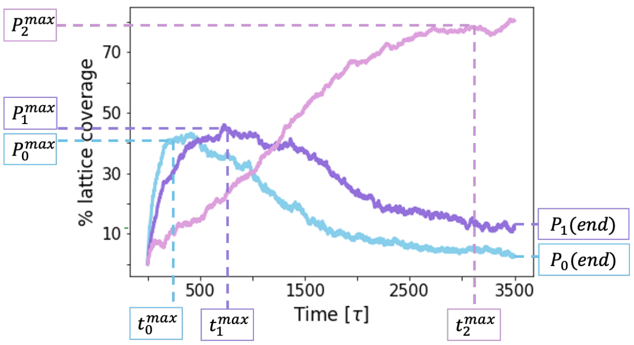


FIG. S2. **Characterising lattice coverage curves.** The Degree of Sequentiality (DoS) quantifies the characteristics of the lattice coverage curves we produce and distinguishes between sequential peaks and non-sequential peaks.

### III. EXPLORING THE VALIDITY OF THE MODEL ASSUMPTIONS

To build the model we had to choose a finite number of sites in the lattice and we made the assumption that the Boltzmann factor in equation (S1) was absorbed by the $r_{\rm off}$ rate. Here we explain the reasoning behind our choices for the lattice length $L$, and for $r_{\rm on}$ and $r_{\rm off}$, and we show that the results are robust against these choices.

The results in the main text were acquired using a lattice of $L = 42$ sites, where each site can host up to 3 monomers of a different type. The number of ESCRT-III monomers in a filament has been estimated at 150 to 300 monomers [17, 25], so that the 126 monomers per filament in our model is of the same order. To check for size effects, we performed a set of simulations where we increased the lattice length up to 5 times, to $L = 210$ sites (630 monomers), while keeping other parameters the same. Fig. S3a shows that the Degree of Sequentiality (DoS) is practically unaffected. The lattice coverage for monomers of different types (in insets) still shows full sequentiality and monomers bind and unbind in order of increasing curvature, as they should. The curves for larger lattices (■) are almost identical to the ones for smaller lattices (▲, which is the same as Fig. 3d). Our results are not affected by the size of the system within physiologically relevant sizes.

Our kinetic Monte Carlo approach conserves the equilibrium Boltzmann distribution. The ratio of the rates with which a certain monomer insertion or removal occurs, i.e. the equilibrium constant of this given process $K_{\rm eq} = r_{\rm on}/r_{\rm off}$, is equal to the exponential of the associated energy change. In our model we choose to assign $r_{\rm on}$ $= 1$ (in units of simulation time steps) and $r_{\rm off} = 1/K_{\rm eq}$. This choice effectively models diffusion-limited binding, whose rate $r_{\rm on}$ depends only on the diffusion coefficient, a constant, and sets the unit of time of our simulations. The unbinding rate $r_{\rm off}$, conversely, depends on the energies associated with the binding into the filament and membrane, through equation (S1).

The exact opposite limit would be to choose $r_{\rm off} = 1$ and $r_{\rm on} = K_{\rm eq}$. All the other choices need to lie in-between. To prove that the results are robust against the choice of the partitioning of $r_{\rm on}$ and $r_{\rm off}$, we repeated our simulations choosing $r_{\rm off} = 1$. While it is not clear which type of a physical scenario this would represent, it is a valid choice within the limits that thermodynamics imposes. Now Eq. (S1), the rate of inserting a monomer of type $i$ at site $m$, is replaced by the following:

$$r_{\rm on}^{m,i} = r_{\rm off}^{i}\; e^{-\beta(-E_i^{\rm adh}+\Delta E_{m,i}^{\rm fil}+\Delta E_{m,i}^{\rm mem})}\,, \tag{S14}$$

where $r_{\rm off}^{i} = 1$ in units of simulation time steps. The resulting curves, shown in Fig. S3b, are qualitatively identical to the ones in Fig. 3d of the manuscript (also reproduced in Fig. S3a ▲). By exploring two diametrically opposite choices, we conclude that the exact expression of the rates will not substantially affect our results.

Finally, it is not surprising that different choices for

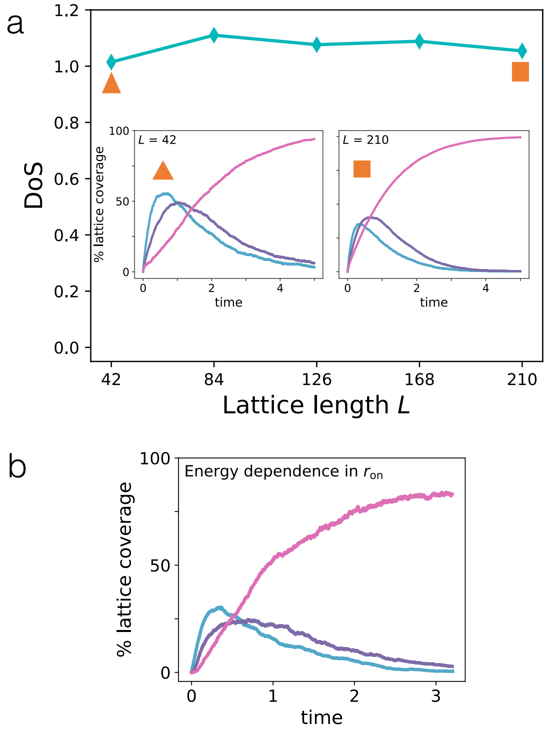


FIG. S3. **Interrogating the model assumptions.** a) The Degree of Sequentiality (DoS), defined in section II, is unaffected by changes in the lattice length, all other parameters staying the same. The insets show Fig. 3d and its equivalent for a 210-site lattice. b) Curves of monomer lattice coverage for a system where the Boltzmann factor is adsorbed in the binding rate, as per Eq. (S14), and the unbinding rates are equal. The sequential peaks are qualitatively unchanged with respect to Fig. 3d, where the dynamics is governed instead by Eq. (S1) and the binding rates are equal.

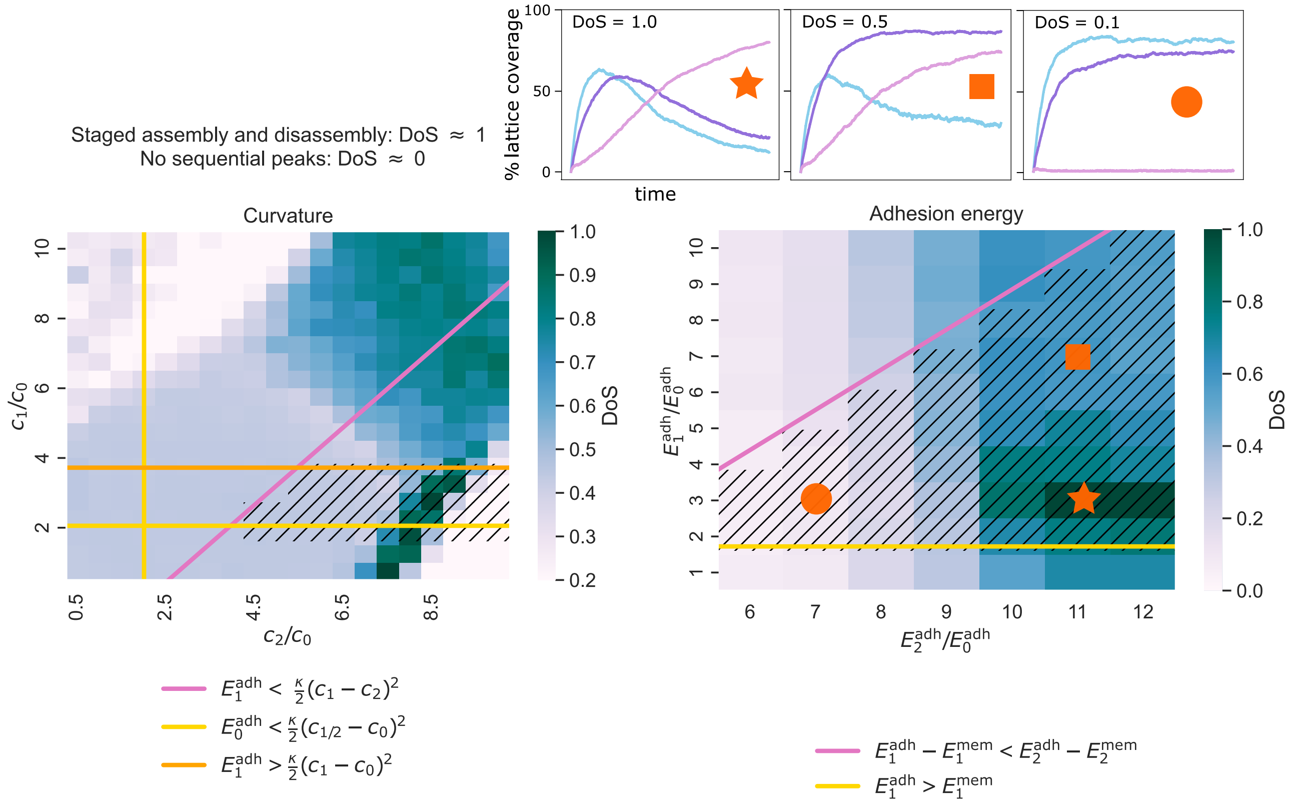


FIG. S4. **Parameter ranges for sequentiality with scaling arguments.** The Degree of Sequentiality (DoS) dependence on properties of monomers 0, 1 and 2. The highest value of DoS (in dark blue) qualitatively gives the best sequential peaks (see panel above b). Monomer 0 is flat ($p_0$=0), while monomers 1 and 2 have pitches $p_1$ = 30nm and $p_2$ = 45nm. Panel a) shows how DoS depends on the curvatures of the high-pitch monomer type (2) and the intermediate-pitch monomer type (1). Adhesion energies are fixed. The strip of high DoS around the bisectrix shows that the curvatures of 1 and 2 can not be too dissimilar. The most sequential peaks are seen for $c_2 > c_1 > c_0$. In b) the adhesion energies of the large-pitch monomer type (2) and the intermediate-pitch monomer type (1) are changed. Sequential peaks are observed when $E_2^{adh} > E_1^{adh} > E_0^{adh}$, in other words, the optimal binding energy ratios were found to be in order of increasing pitch. Both in a) and b), the straight lines show boundaries we can draw from some simple scaling arguments: the hashed regions that they delimit show the parameter space that lies inside these boundaries.

partitioning of energy between $r_{\rm off}$ and $r_{\rm on}$ (while keeping their ratio constant) give the same conclusions, since the two systems feature exactly the same physical ingredients. The different partitioning effectively remaps binding time in a different way for different monomer types. The pronounced free energy differences of our systems overshadow this effect, so that the results are qualitatively unaffected. Our model assumes that the energy barriers for addition of all monomers are comparable, or that they are small compared to the thermodynamic potentials that drive the process. In the absence of more detailed experiments on the kinetics of the transient assemblies, or atomistic molecular dynamics simulations, this is the most fair assumption to make.

### IV. EXPLORING CONDITIONS FOR SEQUENTIAL BINDING/UNBINDING

Having defined an observable (the DoS) that indicates whether a given set of parameters produces sequential peaks in the probability-time plots, we could explore the parameter space via this observable. Using this quantity, we sought to probe how robust the sequential behaviour is in Fig. 3 when varying the parameters, as well as understand this sequential behaviour via energy scaling arguments. The DoS in all our plots was re-normalized using the largest DoS we saw throughout the study (0.187). Defined and separated peaks correspond to a large value of DoS, as in the ★ panel above Fig. S4b, while absence of clear peaks gives a low value of DoS, as in the ● panel above Fig. S4b.

The binding-unbinding dynamics over time of the

monomers in our model is governed by energies, namely adhesion energy, filament bending energy and membrane bending energy. Fig. S4a shows how the DoS varies as the curvatures of monomers 1 and 2 are changed. To ensure that monomer 1 has a preference to bind when monomer 0 has already bound, which happens near the start of the simulation, the adhesion energy of monomers 1 must be larger than the cost of filament bending when monomers 1 and 0 are on the lattice together: $E_1^{\mathrm{adh}} > \frac{\kappa}{2}(c_1 - c_0)^2$. This corresponds to the region below the orange horizontal line in Fig. S4a. However, because monomer 0 has no energy cost due to membrane bending ($p_0 = 0$), and eventually must be forced off the lattice by monomers 1 and 2 binding, the adhesion energy of type 0 must be less than the bending energy of a frustrated filament where monomers of type 0 are bound either with type 1 or type 2: $E_0^{\mathrm{adh}} < \frac{\kappa}{2}(c_1-c_0)^2$ and $E_0^{\mathrm{adh}} < \frac{\kappa}{2}(c_2-c_0)^2$. These two conditions are satisfied in the top right corner of Fig. S4a, as delimited by the two yellow lines. Similarly, monomer 1 needs to be forced off the lattice by monomer 2, so its adhesion energy must be less than the bending energy of a frustrated filament where monomers 1 are bound on a type-2 filament: $E_1^{\mathrm{adh}} < \frac{\kappa}{2}(c_1 - c_2)^2$. This corresponds to the area below the pink line in Fig. S4a. It should be noted that the pink boundary is a generous upper bound for $\frac{\kappa}{2}(c_1-c_2)^2$ as this assumes that the average curvature around a monomer of type 1 is exactly $c_2$ when monomers 1 no longer have a preference to bind, while in actual fact for the sequential peaks to occur the monomers 1 will lose their preference to bind when the average local curvature is somewhere between $c_1$ and $c_2$. The hashed region in Fig. S4a shows the portion of parameter space where all these inequalities hold true. Additionally, the curves also lose the sequentiality characteristic when the curvature of monomer 2 is too large; in this region the adhesion energy of monomer 2 is no longer large enough to overcome the bending energy between monomers 2 and monomers 0/1 and so monomers 2 will not bind. Although membrane bending is not explicitly accounted for in these calculations, the points in parameter space corresponding to the highest DoS fall within the hashed region, suggesting that our model is compatible with such intuitive and qualitative arguments.

Fig. S4b shows how DoS varies as the adhesion energy of monomers 1 and 2 are changed. The monomers with the large pitch, type 2, must be stable once all bound to the lattice, at late times. Hence, the cost of membrane bending per monomer for that type must be smaller than the adhesion energy of that type: $E_2^{\mathrm{adh}} > E_2^{\mathrm{mem}}$, where $E_2^{\mathrm{mem}}$ is the membrane bending energy when only type 2 is present and fills the whole lattice, as per Eqs. (S10) and (S12). This must also be true for monomers 1, which must also stably bind to the lattice at some time: $E_1^{\mathrm{adh}} > E_1^{\mathrm{mem}}$. This condition is met above the yellow horizontal line in Fig. S4b. Lastly, the global energy minimum must feature type 2 dominating on the lattice, therefore the sum of adhesion energy and membrane bending energy per monomer for type 2 has to be more negative than the same quantity for type 1: $-E_2^{\mathrm{adh}} + E_2^{\mathrm{mem}} < -E_1^{\mathrm{adh}} + E_1^{\mathrm{mem}}$. This is true below the pink line in Fig. S4b. Again, the hashed region in Fig. S4b shows the area in parameter space where all these inequalities hold true. It coincides with the area of high DoS from simulations.

### V. EFFECT OF PITCH

Fig. S5 shows the effect of the pitch $p_1$ of type-1 monomers on the binding speed of the monomer types. The binding speed is measured using the time taken for the lattice coverage of a particular monomer type to reach half its maximum lattice coverage, $t_{1/2}$. The smaller $t_{1/2}$, the quicker a monomer type has bound. First of all, the binding of monomers 0 ($p_0 = 0$) is mostly unaffected by $p_1$, as type 0 always binds first and quickest. As for monomers 1, the larger $p_1$, the longer it takes for them to bind. This behaviour is quite intuitive. More interesting is the non-monotonic behaviour in the $t_{1/2}$ of the large-pitch monomer, type 2, which probes the speed of the overall process. The initially decreasing regime is because, as $p_1$ increases, it becomes closer to $p_2$, lowering the energy barrier for binding of type 2. As $p_1$ further increases, though, monomers 2 bind immediately after monomers 1, but monomers 1 take a longer time to bind because their pitch is then further away from $p_0$. In summary, the rate of the binding of type-2 monomers is limited for low $p_1$ by the binding of monomers 2, and for large $p_1$ by the binding of monomers 1. The time for reaching the final state (full binding of type-2 monomers)

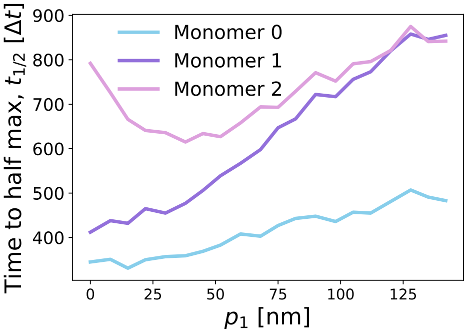


FIG. S5. **The effect of pitch on the binding dynamics.** This plot shows the effect of changing the pitch of monomers 1 on the binding speed of the monomer types. These simulations were all with equal curvatures and binding affinities of the monomers, with no sequential unbinding: $c_0^{-1} = c_1^{-1} = c_2^{-1} = 30\,\mathrm{nm}$, $E_0^{\mathrm{adh}} = E_1^{\mathrm{adh}} = E_2^{\mathrm{adh}} = 6.5 k_{\mathrm{B}}T$, $p_2 = 150\,\mathrm{nm}$, $p_0 = 0$. The quantity $t_{1/2}$ is the time for the lattice coverage of a particular monomer type to reach half its maximum lattice coverage: the faster the monomers bind stably, the smaller the value of $t_{1/2}$.

is minimised for intermediate pitches $p_1$ of the intermediate monomer.

### VI. MEMBRANE-FILAMENT BINDING VERSUS FILAMENT-FILAMENT BINDING.

In our model, adhesion energies do not distinguish between membrane-filament and filament-filament interactions. To understand which interaction is more important for this system, we altered the model so that the adhesion energy had two components, one gained whenever a monomer binds to the lattice, and the other gained when the monomer binds to a lattice site with a monomer already bound. We chose to keep the total possible adhesion energy of each monomer type ($E^{\mathrm{adh}}_{\mathrm{fil-mem}} + E^{\mathrm{adh}}_{\mathrm{fil-fil}}$) the same as in the original model but shifted the weighting of this total adhesion energy between membrane-filament and filament-filament. Fig. S6 shows how the DoS value changes as we change this weighting: as we increase $E^{\mathrm{adh}}_{\mathrm{fil-fil}}$, we decrease $E^{\mathrm{adh}}_{\mathrm{fil-mem}}$ thereby keeping the total adhesion energy constant. It is clear that the important interaction is the membrane-filament interaction. This is because the last filament needs a strong binding to the membrane so it will stay to deform it when the other monomer types have unbound. Otherwise, nothing prevents the last filament from unbinding once the other two have disassembled.

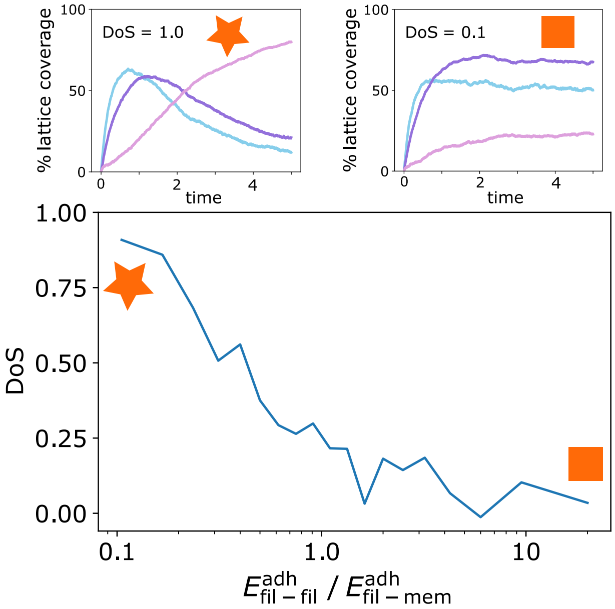


FIG. S6. **Filament-filament versus filament-membrane adhesion energy.** This plot shows how the DoS value changes as we change the weighting of the adhesion energy from more membrane-filament adhesion (left side) to more filament-filament adhesion (right side). The maximum adhesion energy of the monomers is kept the same. The important interaction in our case is the membrane-filament interaction, which is the interaction responsible for a high DoS.

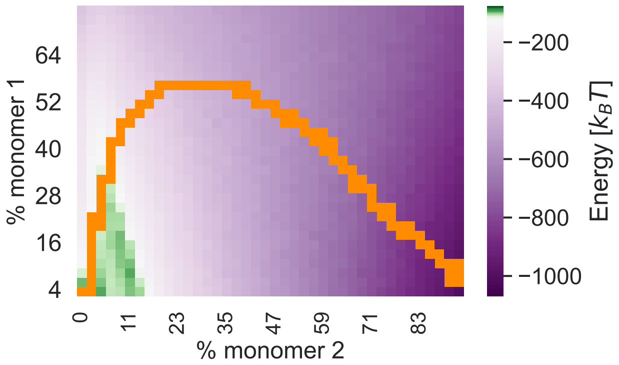


FIG. S7. **Two dimensional energy landscape with sequence pathway.** An energy exploration of number of monomers 1 vs number of monomers 2, including 29% monomers 0 (which is the average number of monomers 0 throughout the sequential pathway). We see an energy barrier (in dark green) to reach the global minimum of full coverage of monomers 2 when not enough monomers 1 are bound. The actual pathway when the full sequence is invoked is shown in orange. The pathway avoids the energy barrier by reaching a minimum number of intermediate monomers, of type 1, on the lattice. The monomers 1 then dissociate and the system reaches eventually its global minimum.

### VII. ENERGY BARRIER IN TWO DIMENSIONS

In Fig. 4a we plotted the average energy while varying the percentage of the highly-curved high-pitch filament (monomer 2). We found that there is an energy barrier to binding these monomers, which is removed when the whole sequence of monomers 0-1-2 is invoked. Fig. S7 shows the full data for when both the number of monomers 2 and number of monomers 1 are changed. The latter shows clearly the energy barrier in achieving the global energy minimum of full type-2 coverage with no monomers 1. The energy barrier (in dark green) extends into the vertical direction, so that there is a minimum number of monomers 1 needed to overcome the barrier. The orange line shows the actual pathway that our system traverses, for parameters that allow full sequentiality and high membrane deformation. It should be noted that Fig. S7 is plotted at constant type-0 coverage (29%, corresponding to the average coverage throughout the time sequence). For the orange pathway, the number of monomers 0 is not constrained.

### VIII. TORSION ENERGY

For simplicity, in our main minimal model we neglect torsional rigidity. This amounts to implicitly assuming that the pitch per turn is much smaller than the radius of

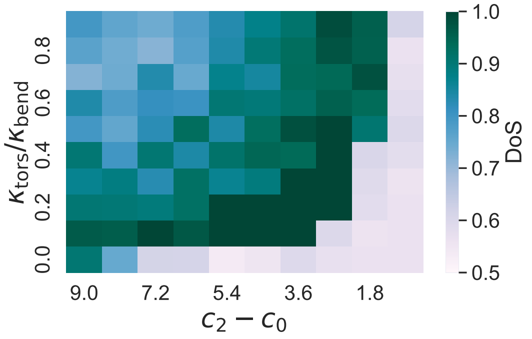


FIG. S8. **Torsional rigidity and curvature differences.** In this plot DoS was calculated while the torsional rigidity $\kappa_{\rm tors}$ was increased from 0 to $\kappa_{\rm bend}$. The difference in curvatures between type-2 and type-0 monomers and also between type-1 and type-0 monomers were changed from $c_2 - c_0 = 9$ down to 0.9 and simultaneously $c_1 - c_0 = 1.5$ down to 0.15. Introducing torsional rigidity has a similar effect as increasing the difference in curvature between the monomers. The boundary of the high-DoS area here shows a $(c_2 - c_0)^2$ dependence, which derives from the energy term in Eq. (S15).

the filaments, and that the torsional rigidity (rigidity of the filaments in the out-of- membrane plane direction) is proportional to the lateral bending rigidity. In this case the bending term of the filament simplifies to only lateral bending and curvature becomes $1/r$. Here we explore the effects on the results when we retract those assumptions.

If $r$ is the radius and $p$ is the pitch per radian, the full equations for curvature $c$ and torsion $\tau$ of an elastic helical rod are as follows:

$$c = \frac{r}{r^2 + p^2}\,,$$
$$\tau = \frac{p}{r^2 + p^2}\,.$$

The torsional energy can then be added to the bending energy to give the following total filament energy, that replaces equation (S3):

$$E^{\rm fil}_{m,i} = \frac{\kappa_{\rm bend}}{2}(\mathcal{C}_m - c_i)^2 + \frac{\kappa_{\rm tors}}{2}(\mathcal{T}_m - \tau_i)^2\,. \qquad \text{(S15)}$$

Here $\mathcal{C}_m$ is the local curvature, $\mathcal{T}_m$ the local torsion, $\kappa_{\rm bend}$ (previously $\kappa$) the bending rigidity and $\kappa_{\rm tors}$ the torsional rigidity. $\mathcal{C}_m$ is mathematically defined in equation (S4), and $\mathcal{T}_m$ is calculated in the equivalent way. We took the filament be of constant length (1000 nm [12]) and kept the total depth (pitch × number of turns) the same as in our original model.

Considering torsion through Eq. (S15) had little to no effect on the results. To get sequential curves with finite torsional rigidity, however, the curvature differences needed to be reduced. This is shown in Fig. S8. Adding torsional rigidity adds another bending term, so to offset its effect and re-establish the sequential curves the curvature differences must be reduced: this makes the filament mechanical energy term once again comparable to the membrane bending term. To conclude, in the model presented in the main text, curvature-induced frustration effectively encompasses the effect of torsional frustration with a re-scaling of the curvature differences.
